# Endothelial CXCL12 is dispensable for adipose tissue function

**DOI:** 10.64898/2026.10.13.758786

**Authors:** Staffan Hildebrand, Max Retiet, Barbara E.F. Pregler, Frederike J. Graelmann, Stefan Löffler, Irmgard Förster, Alexander Pfeifer

**Affiliations:** Institute of Pharmacology and Toxicology, University Hospital Bonn, University of Bonn, Germany; Immunology and Environment, Life and Medical Sciences (LIMES) Institute, University of Bonn, Germany

**Keywords:** Cxcl12, endothelial cells, adipose tissue, thermogenesis, obesity, inflammation

## Abstract

**Background/Objectives:** The adipose tissue endothelium plays a critical role for tissue function in acting as an interface between circulating blood and the various tissue-resident cells. Proper endothelial function ensures, among other things, well-regulated adipose tissue lipid in- and efflux, immune cell diapedesis, and oxygen supply. Dysregulated lipid flux, inflammation, and adipose tissue hypoxia are hallmarks of adipose tissue dysfunction and metaflammation. Additionally, lipid uptake and oxygen supply are important processes enabling heat production in active brown adipose tissue (BAT), which is associated with cardiometabolic health in humans. However, so far, little is known about the role of adipose tissue endothelial cells (ECs) and their intercellular communication with other adipose tissue cell compartments. Particularly, how this communication changes during thermogenic stimuli such as cold exposure is largely unknown.

**Methods:** We used transcriptome analysis of adipose tissue ECs to identify EC *Cxcl12* as a potential regulator of adipose tissue function. *In vitro* experiments were performed on primary sympathetic neurons and immortalized ECs. The role of EC *Cxcl12* in adipose tissue function was analyzed *in vivo* using conditional knock-out mice.

**Results:** *Cxcl12* expression was induced in BAT ECs upon exposure to cold. Recombinant CXCL12 increased axon outgrowth in primary sympathetic neurons and angiogenesis in ECs. Conditional knock-out of *Cxcl12* in ECs did not yield a discernable phenotype in mice either in normal housing or during cold adaption. Additionally, we could show that EC-derived CXCL12 does not play a role in adipose tissue dysfunction during obesity.

**Conclusions:** *Cxcl12* is highly expressed in the adipose tissue endothelium, but appears to be dispensable for adipose tissue function.

## 1. Introduction

Adipose tissue (AT) is the major storage depot for chemical energy in animals. Excessive accumulation of adipose tissue during obesity is associated with a multitude of comorbidities, and is a major contributor to non-communicable disease globally [1]. Moreover, mammalian AT has evolved the capacity to produce heat during cold stress via futile cycling processes, mainly mitochondrial uncoupling in beige and brown adipose tissue (BAT) [2]. This thermogenic activity is of crucial importance for the maintenance of body temperature, especially in small animals with a high surface-to-volume ratio. While human BAT was previously thought to only be important for infants, who are unable to generate heat by shivering, recent studies have clearly demonstrated that also adult humans have recruitable BAT [3–5]. Interestingly, BAT activity in adult humans has been shown to correlate with cardiometabolic health, highlighting the importance of this unique tissue in human health [6].

Both storage of energy and adaptive thermogenesis play critical roles in supporting whole-organism homeostasis in response to shifting environmental variables such as food availability and ambient temperature. As these variables fluctuate, so do the corresponding adaptive processes in AT, including adipocyte specific processes such as hypertrophy, hyperplasia, apoptosis, and “browning” – the conversion of white adipocytes into brown thermogenic adipocytes [7–9]. Additionally, perturbation of environmental variables also influences non-adipocyte specific processes in AT, such as immune cell infiltration, angiogenesis, and sympathetic innervation [10–12].

The vascular endothelium acts as the barrier between the circulation and AT, and therefore plays a critical role in essentially all adaptive processes in AT, including lipid storage and release [13], immune cell diapedesis and inflammation [14], innervation [15], and browning [16]. The vascular endothelium is also highly involved in the chemokine system, both as a source and target of chemokine signaling, as well as by chemokine binding to the endothelial glycocalyx [17,18]. The vascular endothelium is also essential for the adaptive thermogenic remodeling that takes place during chronic cold exposure in BAT [19]. In order to supply the BAT with sufficient oxygen and fuel under conditions of high heat production, cold exposure triggers an angiogenic response in the vascular endothelium, which enhances tissue perfusion [19]. Additionally, nerve bundles in white adipose tissue (WAT) grow in close proximity to the vasculature [20], and ECs have been shown to secrete neurotropic factors that influence peripheral nerve growth [21].

Stromal cell-derived factor 1, or CXC motif chemokine 12 (CXCL12), is an evolutionary conserved and highly ubiquitous chemokine with multiple critical functions in animal physiology [22]. In AT, CXCL12 derived from either smooth muscle cells or adipocytes themselves has been shown to influence tissue homeostasis by regulating sympathetic innervation, brown adipocyte activity, and immune cell recruitment [23–26]. However, the role of endothelial-derived CXCL12 in adipose tissue function has so far not been investigated. Here, we set out to find novel secreted factors derived from the vascular endothelium in AT, that may influence either WAT or BAT function during cold adaptation. By profiling the adaptive response to cold exposure in AT endothelial cells (ATECs) using mRNA sequencing, we found that *Cxcl12* is both highly upregulated in response to cold exposure, and strongly enriched in ATECs compared to other cell types in BAT. However, our data suggest that EC-derived CXCL12 is dispensable for adipose tissue function.

## 2. Materials and Methods

### 2.1. Animals

C57BL/6J mice were obtained from Charles River. Cxcl12^fl/fl^ (B6(FVB)-*Cxcl12^tm1.1Link^*/J) and Cdh5^Cre^ (B6.Cg-Tg(Cdh5-cre)1Spe/J) were crossed to generate Cxcl12^fl/fl^ Cdh5^Cre/+^ (CXCL12-ECKO) and Cxcl12^fl/fl^ Cdh5^+/+^ (WT) mice. Mice were housed with a light and dark cycle of 12 h each and access to chow diet and water *ad libitum*. All animal experiments were approved by the Landesamt für Verbraucherschutz und Ernährung, NRW, Germany, with the approval numbers 81-02.04.2023.A113 and 81-02.04.2015.A202.

### 2.2. Cold exposure

For cold exposure experiments, mice were acclimatized for either 1 week at 18° C or 3 days at 16° C prior to 7 days of exposure to 4° C. The mice were housed in individual cages to prevent mutual warming.

### 2.3. Metabolic characterization

For metabolic characterization of mice, O_2_ consumption and motility were measured at either 23° C or 4° C for 24 h using the Phenomaster system (TSE Systems). During the measurements mice were single caged with access to food and water *ad libitum*. Acute cold exposure of mice was performed for 1 h at 4° C.

### 2.4. Diet-induced obesity

Six-week-old male mice were fed high-fat diet (HFD, Ssniff, D12492) or control diet (Ssniff, D12450B) for 12 weeks. The body weight was monitored weekly. Insulin and glucose tolerance tests were performed week 10 and 11, respectively, and was followed by metabolic characterization and body composition analysis (Bruker Minispec LF50H).

### 2.5. FACS and mRNA sequencing of adipose tissue endothelial cells

Adipose tissues were dissected and digested with Collagenase type 2 to isolate the stromal vascular fraction (SVF). The SVF was enriched for CD31-expressing cells using anti-CD31 microbeads and MACS (Miltenyi) according to the manufacturer’s instruction. The CD31-expressing cells were then labeled with CD31-PECAM PerCP/Cy5.5 (Biolegend, Cat. No. 102420), CD45-LCA clone 30-F11 APC/Cy7 (Biolegend, Cat. No. 103116), CD102-ICAM2 clone 3C4 PE (Thermo Fisher, Cat. No. A15451), CD144-cadherin clone eBioBV13 AF488 (Thermo Fisher, Cat. No. 53-1441-82), and CD146-MCAM clone ME-9F1 BV605 (BD biosciences, Cat. No. 740434). DAPI was used for live/dead staining (Thermo Fisher, Cat. No. D1306). The samples were sorted using a FACSAria III Cell Sorter using FACSDiva software at the Flow Cytometry Core Facility of the University Clinic Bonn (DFG project number 216372401). RNA was isolated using innuSOLV RNA reagent (Analytik Jena, 845-SB-2090100) and sequenced by 3’ mRNA Illumina sequencing at the Next Generation Sequencing Core Facility at the Institute of Human Genetics, Medical Faculty at the University of Bonn.

### 2.6. Migration assay

Functionally immortalized murine microvascular endothelial cells (CI-muMECs, InSCREENex GmbH, Cat. No INS-CI-1004) were starved for 24 hours to prevent proliferation, and seeded in 24-well inserts with 8 µm pore size (QCM 24-Well Colorimetric Cell Migration Assay, Merck, Cat. No. ECM 508) and incubated for 8 hours. Quantification of migrated cells proceeded according to the manufacturer’s instructions.

### 2.7. Proliferation assay

CI-muMECs were assayed using Click-iT® EdU Alexa Fluor® 488 Imaging Kit (Invitrogen, Cat. No. C10337). Cells were seeded in a µ-Slide 8 well (Ibidi GmbH) and incubated for 16 hours in the presence of EdU and assayed according to the manufacturer’s instructions.

### 2.8. Tube formation assay

CI-muMECs were seeded on 60 µl solidified Matrigel® (Corning, Cat. No. 356234) at a density of 14000 cells per 96-well. CXCL12 (100ng/ml) was added and the plate was incubated for 3 hours. The total tube lengths were measured using the straight line tool in ImageJ (Schneider et al., 2012).

### 2.9. Stromal vascular fraction spheroids

BAT tissues were dissected and digested with Collagenase type 2 to isolate the SVF. SVF cells were seeded in 6-wells in a 50:50 mixture of DMEM with 10% FCS and CI-muMEC growth medium (InSCREENex GmbH). The entire SVF fraction from both intrascapular BAT lobes were seeded in each 6-well. At 90% confluence, the cells were detached and seeded in low-attachment round-bottom 96-well plates at a density of 50000 cells per well. When spheroids were formed, they were transferred to a new 96-well plate with 40µl ECM-Gel (Merck, Cat. No. E1270) per well. After two days, adipogenic differentiation was induced with 500 µM 3-Isobutyl-1-methylxanthin, 1 µM Dexamethasone, 20 nM T3, and 1 nM Insulin, in the presence or absence of CXCL12 (100 ng/ml). Two days after induction, the induction medium was removed and replaced with differentiation medium (DMEM with 10% FCS, 20 nM T3, and 1 nM Insulin) with or without CXCL12. Spheroids were harvested on day 7 after induction, fixed with 4% paraformaldehyde, and stained with anti-CD31 (R&D Systems, Cat. No. AF3628) and anti-Perilipin (Cell Signaling, Cat. No. 9349) antibodies. Afterwards, the stained spheroids were cleared using benzyl alcohol and benzyl benzoate (BABB).

### 2.10. Primary sympathetic neurons

The superior cervical ganglia from newborn mice were isolated and digested with Collagenase IV. The resulting cell suspension was plated on 6-well plates for one hour to allow non-neuronal cell to attach. Non-attached cells were then collected and seeded on poly-D-lysin/laminin-coated coverslips in DMEM with 10% FCS and NGF (100 ng/ml). Cells were allowed to attach and grow for six hours in the presence or absence of CXCL12 (100 ng/ml) before fixation and staining with anti-β3-Tubulin (Abcam, Cat. No. ab18207).

### 2.11. Transverse thick-section brown adipose tissue whole-mounts

Whole BAT tissues were fixed overnight in 4% paraformaldehyde, and 500 µm thick transverse sections were prepared using a vibratome. The sections were then incubated with the primary antibodies (anti-CD31; R&D Systems, Cat. No. AF3628 and β3-Tubulin; Abcam, Cat. No. ab18207) overnight. After washing, the sections were incubated fluorescently labelled secondary antibodies (Alexa Fluor®, Thermo Fischer) overnight, followed by washing and clearing in BABB. Images were taken with a Zeiss AXIO Imager.M2m equipped with Apotome 3. Z-stacked tile-scans were assembled to generate an image of the full transverse section at a Z-depth of approximately 30µm.

### 2.12. FACS-analysis of adipose tissue immune cells

For flow cytometry analysis of immune cell populations in BAT, WATg, and WATi SVF, a piece of tissue was excised, weighed, and minced into small pieces. For digestion, the organ pieces were incubated in a digestion mix of DMEM containing 3 mg/mL BSA and 2 mg/mL Collagenase IV for 30-45 minutes at 37° C upon regular shaking. The digestion was stopped with 20% FCS in PBS, and the samples were centrifuged for 5 minutes. All centrifugation steps were performed at 200g and 4° C, if not stated otherwise. Afterwards, the floating fraction of mature adipocytes was removed, and the cell suspension was filtered through a 70 µm filter. Subsequently, after 10 minutes of centrifugation, red blood cell lysis was performed using 1x RBS Lysis buffer in distilled water for 3 minutes on ice. The cells were centrifuged, resuspended in FACS Buffer (2 mM EDTA and 1% FCS in PBS), and transferred into a 96-well plate. After centrifugation, the cell pellet was resuspended in FACS buffer containing 1% Fc block (anti CD16/32) and 4% rat serum for 20 minutes at 4° C, followed by a surface marker staining for 30 minutes at 4° C with CD11b BUV395 (BD, Cat. No. 563553), Ly6G BUV563 (BD, Cat. No. 612921), CD19 BUV737 (BD, Cat. No. 612781), NK1.1 BV421 (BioLegend, Cat. No. 108731), F4/80 BV510 (BD, 569615), TCRb BV605 (BioLegend, Cat. No. 109241), Ly6C BV711 (BioLegend, Cat. No. 128037), CD11c FITC (BioLegend, Cat. No. 117306), CD45 PE (BioLegend, Cat. No. 103106), Siglec-F/CD170 PE Dazzle (BioLegend, 155530), MHCII/ I-A/I-E PE-Cy7 (BioLegend, Cat. No. 107630), and CD206/ Mannose R (BioLegend, Cat. No. 141708). Fixable Viability dye eFluor780 (eBioscience, Cat. No. 65-0865-18) was used for live/dead staining. After washing the stained samples with FACS buffer, data were recorded using the FACSymphony A5 (BD BioSciences) and analyzed in FlowJo Software.

To exclude debris, samples were gated based on FSC-H/SSC-H, and subsequently, single cells were selected using FSC-A/FSC-H and SSC-A/SSC-H. Dead and CD45-cells were excluded before further gating based on antibody staining to identify target cell populations. B cells were identified as CD19+ cells, whereas NK Cells were characterized as NK1.1+ cells and T cells were TCRb+. Neutrophils were identified as CD11b+ Ly6G+ and eosinophils as CD11b+ Siglec-F+ cells. Macrophages were identified as CD11b+ F4/80+, and macrophage subtypes were distinguished by their expression of CD11c and CD206. DCs were gated as CD11b+/- CD11c+ MHCII+ cells, and monocytes were identified as CD11b+ Ly6C+ MHC2-cells.

### 2.13. RT-qPCR

Total RNA was isolated with innuSOLV RNA reagent (Analytik Jena, 845-SB-2090100), and reverse-transcribed using cDNA synthesis kit (ProtoScript II First Strand cDNA Synthesis Kit, NEB). Quantitative PCR was performed using the cDNA as template with Luna® Universal qPCR Master Mix (NEB). *Hprt* was used as housekeeping gene.

### 2.14. Western blot

Cells were lysed with radioimmunoprecipitation assay (RIPA) buffer containing protease and phosphatase inhibitors. SDS-PAGE was performed in SDS-containing electrophoresis buffer at room temperature and proteins were transferred to PVDF membranes using the Mini Trans-Blot® Cell system (Bio-Rad Laboratories, Hercules, USA). After transfer, the membrane was blocked in TBS with 0.1% Tween-20 (TBST) and 5% milk protein (AppliChem GmbH, Darmstadt, Germany) for 1 h. Membranes were incubated with primary antibody at 4° C overnight, washed, and incubated with secondary HRP-conjugated antibody for 1 h at room temperature. The membrane was then washed and covered in ECL reagent for 2 min according to the manufacturer’s instructions. Luminescence was measured immediately in an ImageQuant LAS 4000 chemiluminescence reader. Densitometric analysis was performed in ImageJ.

### 2.15. Histology

Adipose tissues were fixed in PBS containing 4% PFA overnight and dehydrated using increasing ethanol concentrations. The paraffin-embedded tissue was cut into 5 μM sections. For H&E staining, sections were stained with haematoxylin (Merck, 1.09249) for 2 min and washed with deionized water, followed by staining with eosin (Merck, 1.09844) by incubating sections in eosin for 1 min and then washing for 4 min with deionized water.

### 2.16. Analysis of single nucleus sequencing data from human PVAT

Single cell sequencing data from brown adipose tissue was downloaded from the Gene Expression Omnibus, accession number GSE160585 [27]. Single cell sequencing data from white adipose tissue was downloaded from the Single Cell Portal (https://singlecell.broadinstitute.org/single_cell), study number SCP1376. Data was processed using the Seurat package.

### 2.17. Statistical analysis

Differences between two independent samples were evaluated using two-tailed Student’s t-tests, unless otherwise stated.

## 3. Results

We first isolated ATECs from BAT, inguinal (subcutaneous) WAT (WATi), and gonadal (visceral) WAT (WATg), of either cold-exposed mice or mice housed at room temperature. Enrichment of the stromal vascular fraction (SVF) for ECs expressing the pan-EC marker CD31 *via* Magnet-assisted cell sorting (MACS) was followed by Fluorescence-activated cell sorting (FACS), using antibodies for the EC markers CD31, CD102, and CD146. The immune cell maker CD45 was included in the dump channel to exclude potentially co-enriched immune cells. All CD45^-^ CD31^+^ cells also expressed CD102 and CD146, suggesting a highly pure population of ECs (Supplemental Figure 1). Isolated CD45^-^ CD31^+^ CD102^+^ CD146^+^ ECs from the adipose tissues of mice housed at either 23° C or 4° C for one week were then analyzed by 3’ mRNA sequencing (Figure 1A). Principal component analysis showed clear phenotypic differences of ECs from each depot, as well as a distinct effect of cold exposure on the transcriptome of BAT-ECs (Figure 1B). ECs from all depots and conditions showed high expression of typical EC markers such as *Gpihbp1*, *Pecam1*, and *Cdh5*, while showing low expression of immune cell (*Adgre1, Ptprc*), preadipocyte and pericyte (*Pdgfra, Pdgfrb*), and mature adipocyte (*Cidea, Ucp1, Plin, Adipoq, Lep*) markers, demonstrating that the isolation method used generated a pure population of ECs (Figure 1C). Furthermore, clustering analysis showed a clear impact of cold exposure on the ATEC transcriptome, particularly in the BAT-ECs (Figure 1D). In order to identify cold-regulated secreted factors that may mediate EC-BA crosstalk during thermogenic activation, we filtered the differentially expressed genes from BAT-ECs on Uniprot annotation, focusing only on genes coding for proteins that are annotated as secreted. This analysis yielded several candidates that were significantly upregulated in the BAT-ECs of cold-exposed mice, with *Cxcl12* showing the most significant upregulation (Figure 1E). Comparison of read counts of *Cxcl12* in ECs across all AT depots and conditions showed that *Cxcl12* was highly expressed basally in all depots, but was only induced by cold in thermogenically active ATECs (BAT and WATi, Figure 1F). Re-analysis of previously published single-cell sequencing of BAT SVF [27] confirmed that *Cxcl12* is highly expressed in BAT-ECs, and is more highly expressed in ECs than other BAT SVF cell types (Figure 1G).

**Figure 1.**
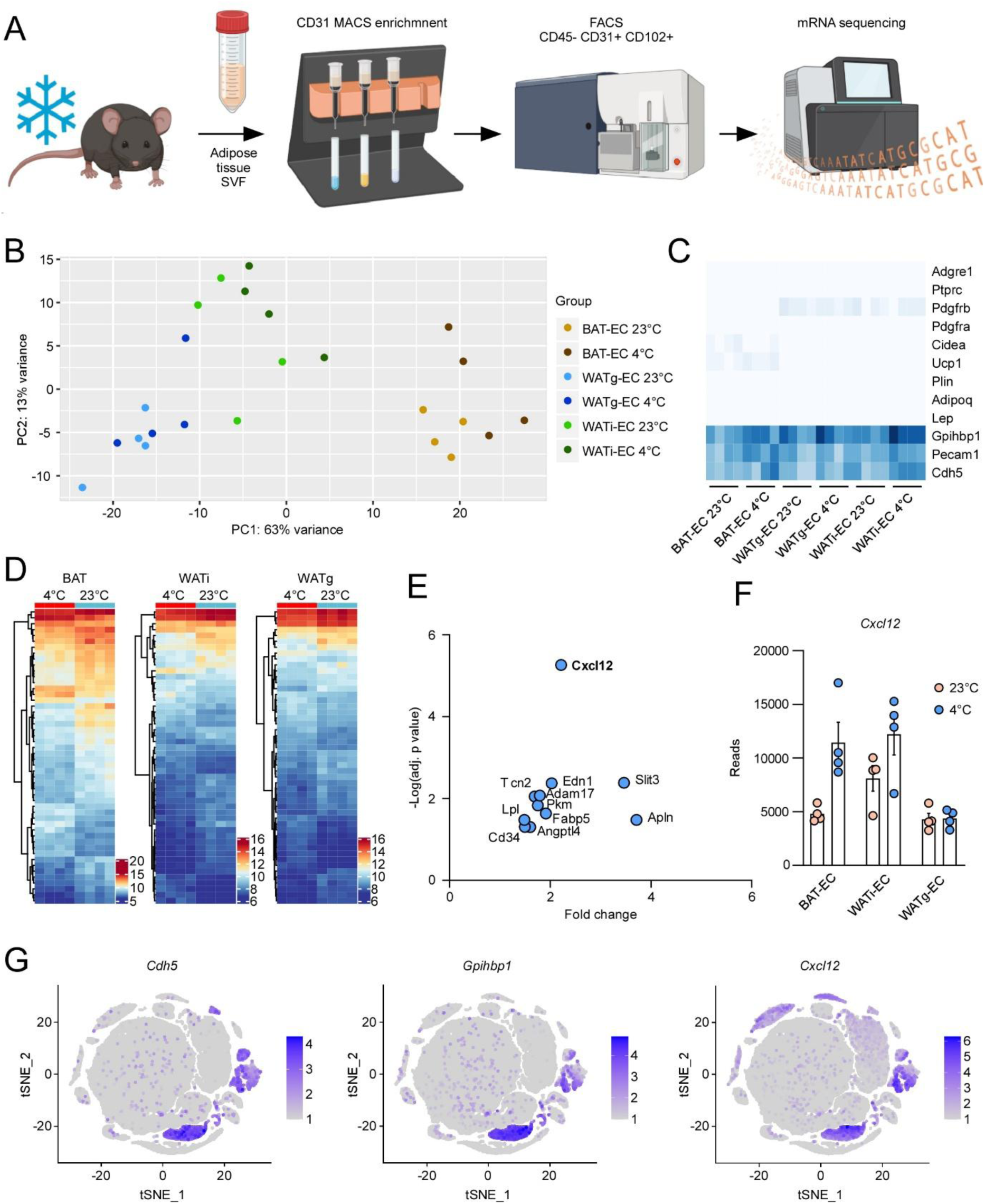
Analysis of the adipose tissue EC transcriptome during cold adaptation. (**A**) Scheme depicting the experimental pipeline. (**B**) Principal component analysis of the EC transcriptome from BAT, WATi, and WATg in cold-exposed or control mice. (**C**) Heatmap showing typical macrophage, perivascular, adipocyte, and EC marker expression in the adipose tissue ECs of cold-exposed or control mice. (**D**) Clustering analysis showing transcriptome changes induced by cold exposure in ECs from different adipose tissues. (**E**) Scatter plot showing the upregulated genes encoding secreted factors in BAT ECs. (**F**) Expression of *Cxcl12* in different adipose tissue ECs at 23° C and 4° C. (**G**) Single-cell expression of *Cdh5*, *Gpihbp1*, and *Cxcl12* in adipose tissue SVF (data retrieved from Gene Expression Omnibus entry GSE160585 [27]).

CXCL12 has previously been shown to regulate angiogenesis in ECs derived from human umbilical veins (HUVEC) [28]. To see if CXCL12 also regulates endothelial cell function in the microvasculature, we treated murine microvascular endothelial cells (muMECs) with 100 ng/ml CXCL12 and measured their angiogenic response. Treatment with CXCL12 significantly increased transwell migration, tube formation on Matrigel, as well as proliferation (Figure 2A-C), suggesting an important role of CXCL12 on microvascular angiogenesis. Next, we investigated the effect of CXCL12 on native BAT ECs by generating murine BAT SVF spheroids in extracellular matrix (ECM) gel. Treatment of BAT SVF spheroids with CXCL12 led to increased sprouting in undifferentiated spheroids, and immunostaining for CD31 revealed significantly increased spheroid vascularization at the end of differentiation, confirming our results in muMECs (Figure 2D-E). Additionally, several reports have shown that CXCL12 increases neurite outgrowth in neurons of the central nervous system [29,30]. However, if CXCL12 can regulate neurite outgrowth in the peripheral postganglionic neurons that innervate BAT (i.e. peripheral sympathetic neurons) is currently unknown. An increased sympathetic innervation of BAT could potentially result in an increased total NE output during BAT activation and therefore increased energy expenditure. We found that treatment of primary murine sympathetic neurons, isolated from the superior cervical ganglia, with 100 ng/ml CXCL12 significantly enhanced neurite outgrowth, with a significantly increased number of total and terminal branches after only six hours of treatment (Figure 2F). Taken together, these data suggest a role for CXCL12 in thermogenic adaptation and remodeling of BAT, by increasing vascularization and sympathetic innervation.

**Figure 2.**
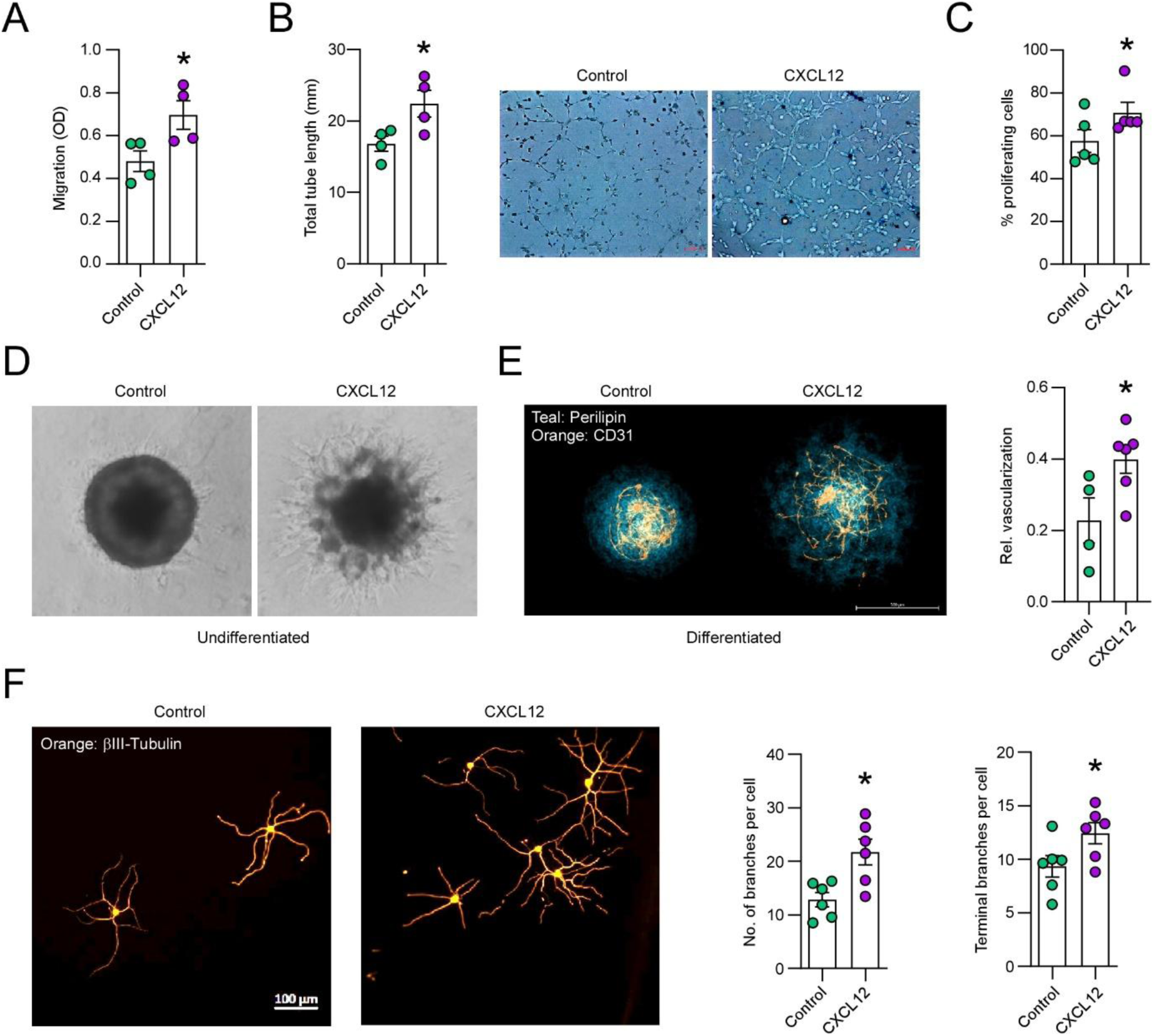
Effect of exogenous CXCL12 on microvascular angiogenesis and sympathetic neurite outgrowth. (**A**) Migration of muMECs in response to 100 ng/ml CXCL12. **(B**) Tube formation in muMECs treated with 100 ng/ml CXCL12. (**C**) Proliferation of muMECs measured by EdU incorporation in response to 100 ng/ml CXCL12. (**D**) Sprouting of undifferentiated BAT SVF spheroids in Matrigel in response to 100 ng/ml CXCL12. (**E**) Vascularization analysis in differentiated BAT SVF spheroids treated with 100 ng/ml CXCL12. (**F**) Neurite outgrowth in primary sympathetic SCG neurons treated with 100 ng/ml CXCL12. *: p<0.05. Student’s t-test.

To validate these results *in vivo*, we generated mice that lack CXCL12 in ECs by crossing Cxcl12^fl/fl^ mice with Cdh5^Cre/+^ mice (CXCL12-ECKO). We validated the *Cxcl12* knock-out by MACS-sorting CD31-expressing ECs from the BAT SVF of CXCL12-ECKO and WT littermates, and found an almost complete absence of *Cxcl12* expression in the CD31^+^ ECs of CXCL21-ECKO mice, but not in the CD31^-^ fractions (Figure 3A). Interestingly, *Cxcl12* expression was also reduced in the floating cell fraction, which is often considered to be composed on only mature (lipid-filled) adipocytes. However, this fraction also expressed relatively high levels of *Pecam1*, suggesting that tissue disruption and Collagenase digestion is insufficient for complete separation of ECs from the mature adipocytes, as has been previously suggested [31]. Nevertheless, *Pecam1* expression was the highest in the CD31^+^ fraction, which also had a near-complete reduction in *Cxcl12*, demonstrating efficient knock-down of *Cxcl12* in the ECs of Cxcl12^fl/fl^ Cdh5^Cre/+^ mice (Figure 3A).

**Figure 3.**
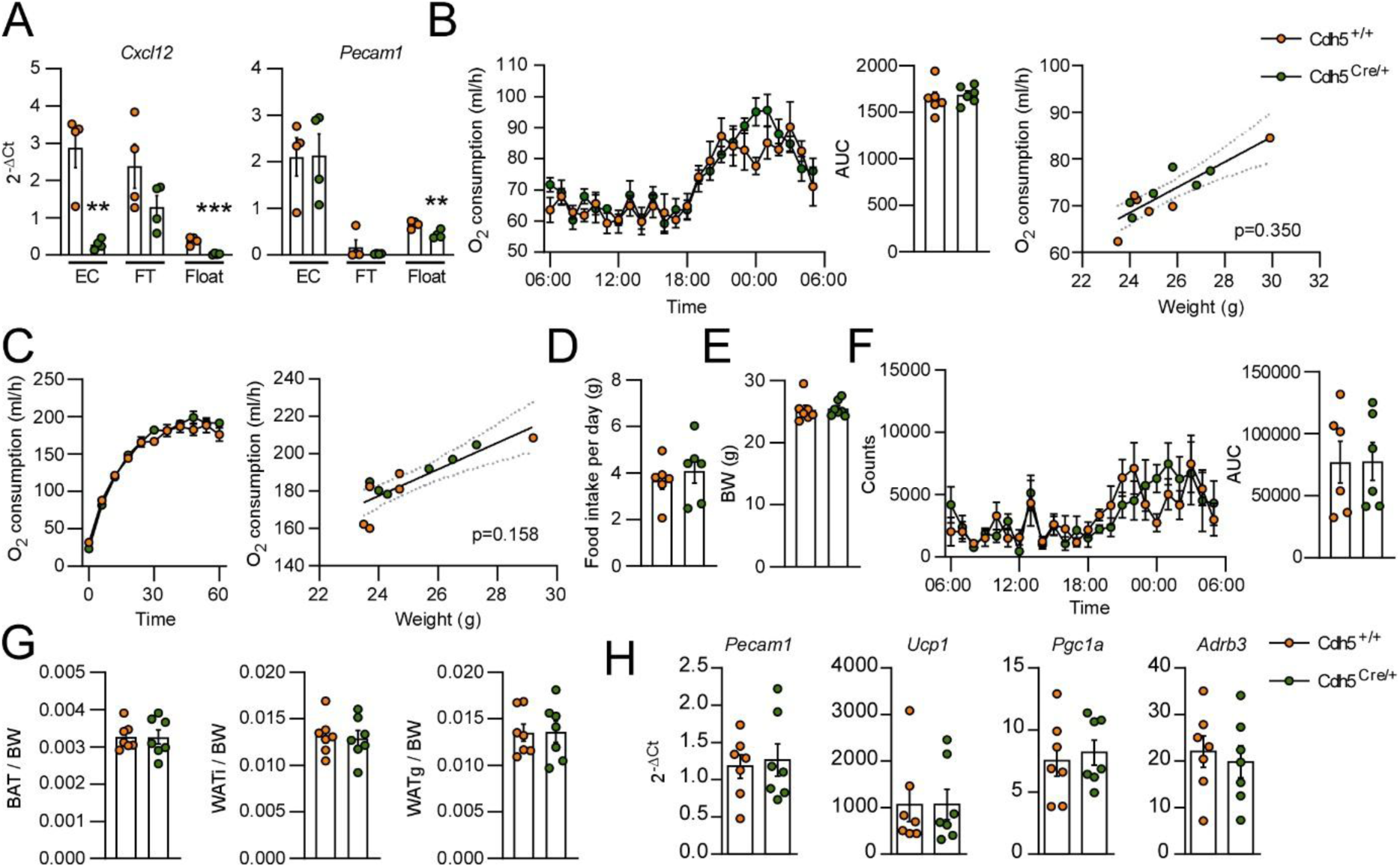
Metabolic characterization of CXCL12-ECKO (Cdh5^Cre/+^) mice under normal housing conditions. (**A**) Expression of *Cxcl12* and *Pecam1* in different BAT cellular fractions following CD31-based MACS enrichment (Student’s t-test). FT: flow-through, Float: floating fraction. (**B**) O_2_ consumption of CXCL12-ECKO mice and WT (Cdh5^+/+^) littermates under normal housing conditions (RT) (ANCOVA). (**C**) O_2_ consumption of CXCL12 mice and WT littermates during a 1-hour cold (4° C) challenge (ANCOVA). (D) Daily food intake of CXCL12-ECKO mice and WT littermates. (**E**) Body weight (BW) of CXCL12-ECKO mice and WT littermates. (**F**) 24-hour motility of CXCL12-ECKO mice and WT littermates housed at RT. (**G**) BAT, WATi, and WATg mass in relation to BW of CXCL12-ECKO mice and WT littermates. (**H**) mRNA expression of *Pecam1*, *Ucp1*, *Pgc1a*, and *Adrb3* in BAT of CXCL12-ECKO mice and WT littermates. **: p<0.01, ***: p<0.001.

Next, we subjected CXCL12-ECKO and WT littermates to rigorous metabolic phenotyping. Surprisingly, O_2_ consumption measurement of CXCL12-ECKO mice in metabolic cages did not reveal any significant differences in energy expenditure in mice housed at 23° C (Figure 3B). Additionally, CXCL12-ECKO mice challenged with an acute (1 hour) bout of cold exposure did not display any difference in thermogenic capacity compared to their WT littermates, as measured by peak acute cold-induced O_2_ consumption (Figure 3C). Food intake, as well as body weight, 24-hour motility, and adipose tissue mass was also unchanged in CXCL12-ECKO mice (Figure 3D-G). Furthermore, the expression of the typical thermogenic genes *Ucp1*, *Pgc1a*, and *Adrb3* were not significantly dysregulated in the BAT of CXCL12-ECKO mice (Figure 3H). Expression of the pan-endothelial marker *Pecam1* was also unchanged, suggesting that vascularization is normal in CXCL12-ECKO mice (Figure 3H). Similarly, thermogenic, adipogenic, and endothelial markers were unchanged in WATi and WATg of CXCL12-ECKO mice (Supplemental Figure 2A-B). Histological examination by H&E staining of BAT, WATi, and WATg sections did not reveal any differences in the gross tissue morphology of the adipose tissues of CXCL12-ECKO mice compared to their WT littermates (Supplemental Figure 2C).

Taken together, these data indicate that EC-derived CXCL12 is dispensable for BAT function in mice under normal (room temperature, RT) housing conditions.

Next, we investigated the effect of endothelial CXCL12 during chronic (one week) cold exposure. As our *in vitro* experiments suggested an important role of CXCL12 for both BAT EC angiogenesis and sympathetic innervation, we expected a defect in cold adaptation in mice lacking EC-derived CXCL12. However, we could not detect any functional difference in thermogenic activity of the CXCL12-ECKO mice during cold adaption, compared to WT littermates. CXCL12-ECKO mice also displayed normal energy expenditure, body weight, food intake, adipose tissue mass, and motility (Figure 4A-E). To investigate whether the CXCL12-ECKO mice had reduced sympathetic innervation in the BAT, we performed Western blot analysis of tyrosine hydroxylase (TH), the rate-limiting enzyme in norepinephrine synthesis in peripheral sympathetic neurons. Interestingly, we could not detect any difference in TH expression in the BAT of CXCL12-ECKO after chronic cold exposure (Figure 4F). Previous reports have suggested that CXCL12 may also be expressed in mature brown adipocytes [24]. However, *Cxcl12* expression was almost completely absent in whole BAT lysates of CXCL12-KO mice, suggesting that ECs are the major source of CXCL12 in BAT (Figure 4G). This is also corroborated by the single-cell sequencing and qPCR data from MACS-sorted SVF shown above (Figure 1G and 3A, respectively). Despite the strong knock-down of *Cxcl12*, no effect could be seen in the expression of either endothelial (*Pecam1*) or thermogenic (*Ucp1*, *Pgc1a*, *Adrb3*) marker expression in the BAT of CXCL12-ECKO mice (Figure 4G). Additionally, no difference could be seen in the tissue and cellular morphology in either BAT or WAT of cold-exposed CXCL12-ECKO mice (Figure 4H-I). These data strongly suggest that endothelium-derived CXCL12 is dispensable for BAT thermogenic capacity and adaptation to cold.

**Figure 4.**
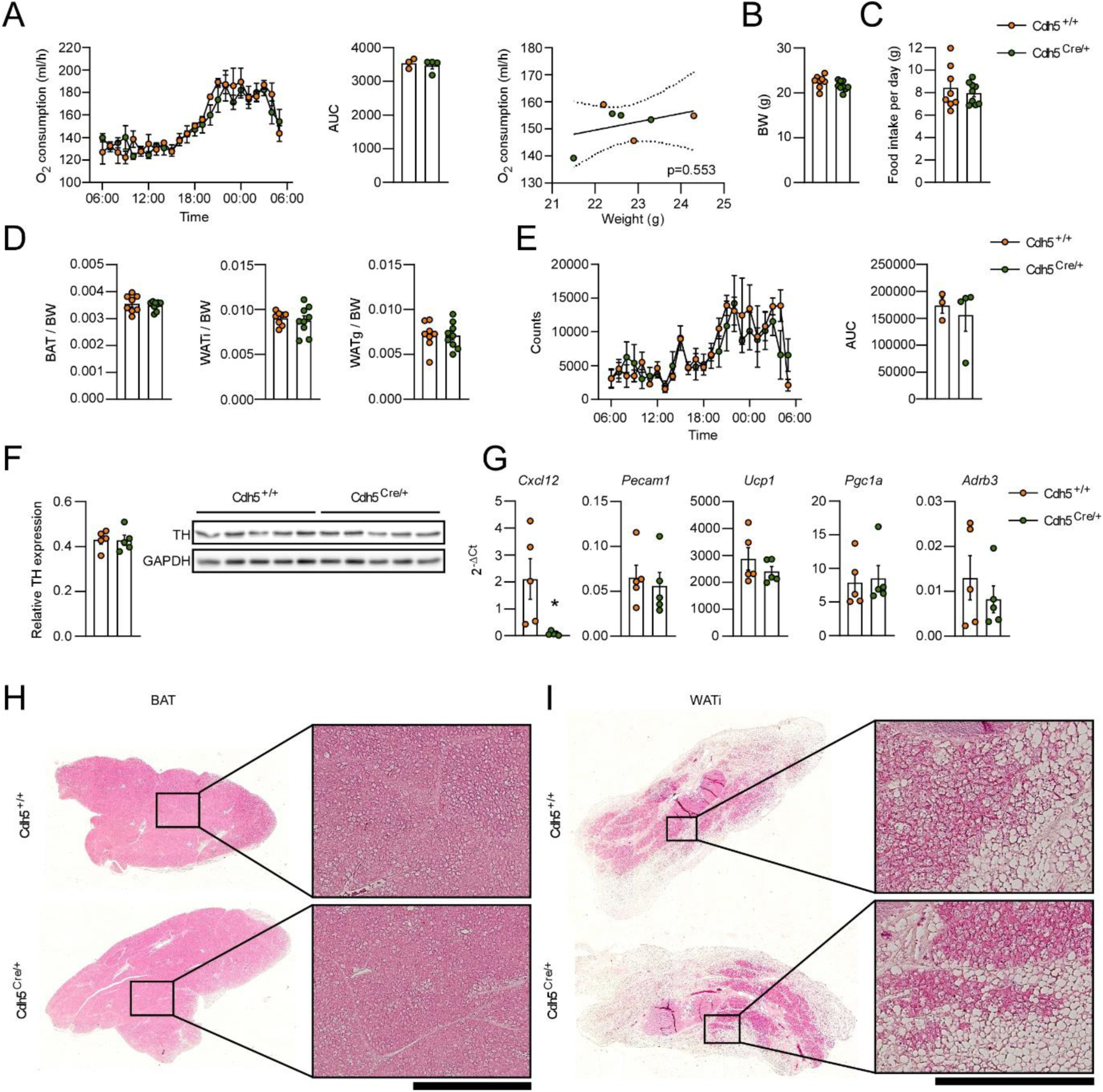
Metabolic characterization of CXCL12-ECKO mice during cold adaptation. (**A**) O_2_ consumption of CXCL12-ECKO mice and WT (Cdh5^+/+^) littermates housed at 4° C for one week (ANCOVA). (**B**) Body weight (BW) of CXCL12-ECKO mice and WT littermates. (**C**) Daily food intake of CXCL12-ECKO mice and WT littermates. (**D**) BAT, WATi, and WATg mass in relation to BW of CXCL12-ECKO mice and WT littermates. (E) 24-hour motility of CXCL12-ECKO mice and WT littermates housed at 4° C for one week. (**F**) Western blot analysis showing tyrosine hydroxylase (TH) expression in the BAT of cold-exposed CXCL12-ECKO mice and WT littermates. (**G**) mRNA expression of *Cxcl12*, *Pecam1*, *Ucp1*, *Pgc1a*, and *Adrb3* in BAT of CXCL12-ECKO mice and WT littermates. *: p<0.05. Student’s t-test. (**H**) H&E staining of histological sections from BAT of CXCL12-ECKO mice and WT littermates. Scale bar = 500 µm. (**I**) H&E staining of histological sections from WATi of CXCL12-ECKO mice and WT littermates. Scale bar = 500 µm.

Given the strong effect of CXCL12 on neurite outgrowth seen in primary sympathetic neurons *in vitro*, the lack of phenotype of CXCL12-ECKO mice is surprising, especially given the known proximity of nerve bundles and the vasculature in WAT. To further investigate the interactions between peripheral neurons and the vasculature in BAT, we performed immunostaining for endothelial cells and neurons in a cleared transverse whole-mount section of BAT. As has previously been shown in WAT [20], we could identify large nerve bundles in close proximity to the larger vessels (Figure 5A, inset 1). Surprisingly, however, we could not see any co-localization of finer neurites with the microvasculature (Figure 5A, inset 2). Indeed, detailed analysis of *Cxcl12* expression in BAT-ECs revealed a correlation of *Cxcl12* expression with *Rbp7*, which has previously been identified as a marker for large capillaries, as well as with *Car4*, a capillary marker [32]. Importantly, *Cxcl12* was poorly correlated with markers of larger arteries and veins, such as *Mgp*, *Vwf*, and *Cfh* (Figure 5B). This relatively low expression of *Cxcl12* in the ECs from larger vessels that are in closer proximity to nerve bundles, compared to microvascular ECs, may explain the lack of thermogenic phenotype in CXCL12-ECKO mice, despite the clear effect of CXCL12 on neurite outgrowth *in vitro*.

**Figure 5.**
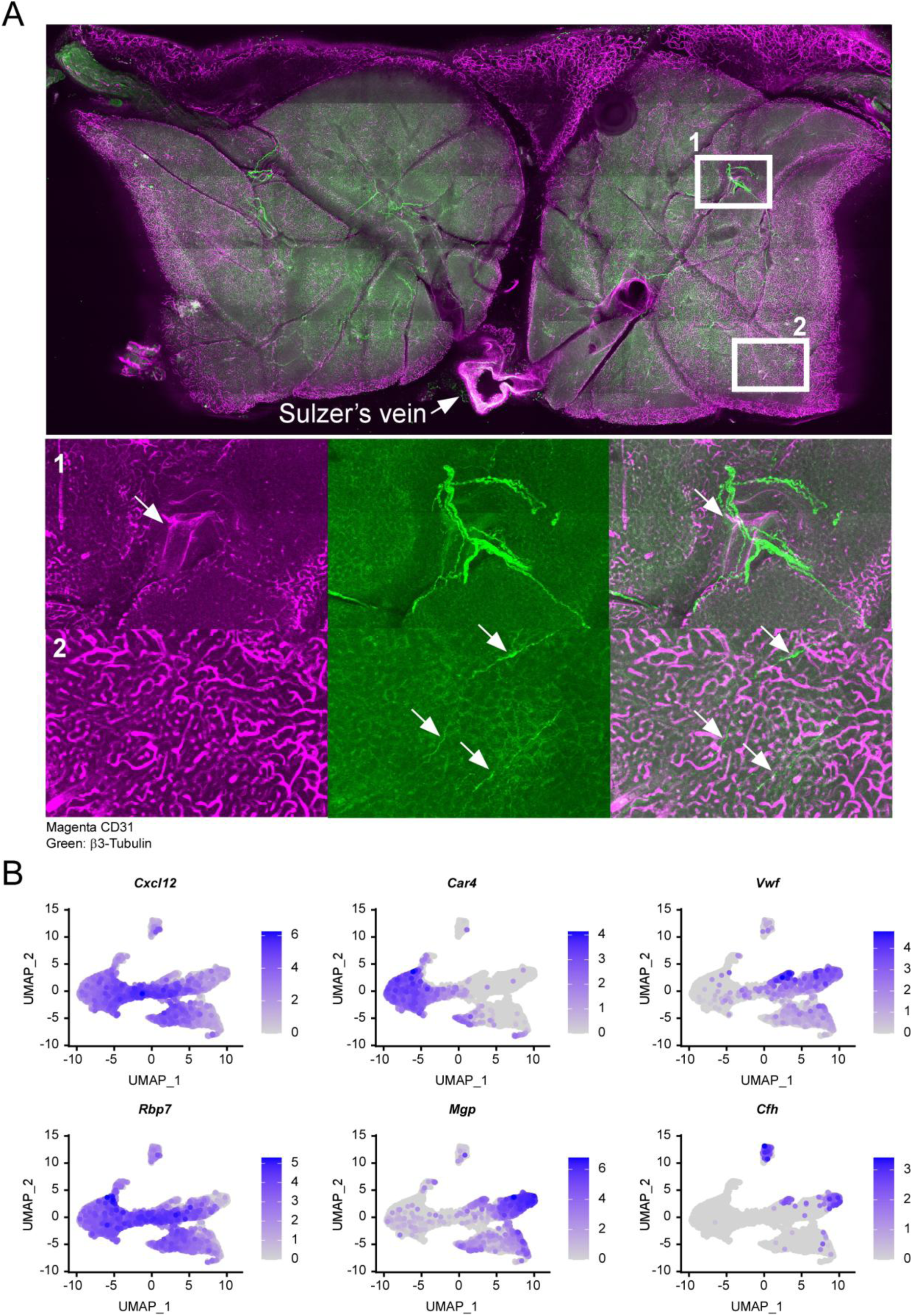
Fine neurites do not associate closely with the microvasculature in BAT. (**A**) Maximum intensity projection of tile-scanned Z-stacks of a cleared transverse whole-mount section of BAT stained with antibodies for CD31 (magenta) and β3-Tubulin (green). (**B**) Analysis of single-cell sequencing data of BAT-EC showing *Cxcl12* expression primarily in large capillaries (data retrieved from Gene Expression Omnibus entry GSE160585 [27]).

Interestingly, analysis of *Cxcl12* expression in the ECs of WATi revealed a close correlation of *Cxcl12* expression with the expression of genes crucial for EC lipid transport (e.g. *Gpihbp1* and *CD36*), as well as immune cell diapedesis (e.g. *Cd300lg* and *Sparc*) (Figure 6A-B). CXCL12 has previously been shown to not only act as a homeostatic chemokine, but also as a chemoattractant during inflammation, potentially by binding to endothelial glycosaminoglycans for presentation to passing immune cells [33–35]. Therefore, we postulated that CXCL12 may be important for either lipid handling or inflammation in WAT. To test this hypothesis, we fed CXCL12-ECKO mice and WT littermates a high-fat diet (HFD) for 12 weeks as a model for diet-induced obesity (DIO). Similarly to metabolically healthy mice, CXCL12-ECKO mice fed a HFD did not show differences in energy expenditure either under normal housing temperature, or during an acute (1 hour) cold challenge (Figure 6C-D). Body weight gain, insulin tolerance, glucose tolerance, and body composition also remained unaltered by EC-specific CXCL12 deletion (Figure 6E-H). RT-qPCR analysis of the different adipose tissue depots of CXCL12-ECKO mice and their WT littermates likewise did not show any difference in *Ucp1* expression or other adipogenic, thermogenic, or vascular markers (Supplemental Figure 3A-C). 24-hour motility and food consumption also remained unchanged in CXCL12-ECKO mice compared to their WT littermates (Supplemental Figure 3D-E).

**Figure 6.**
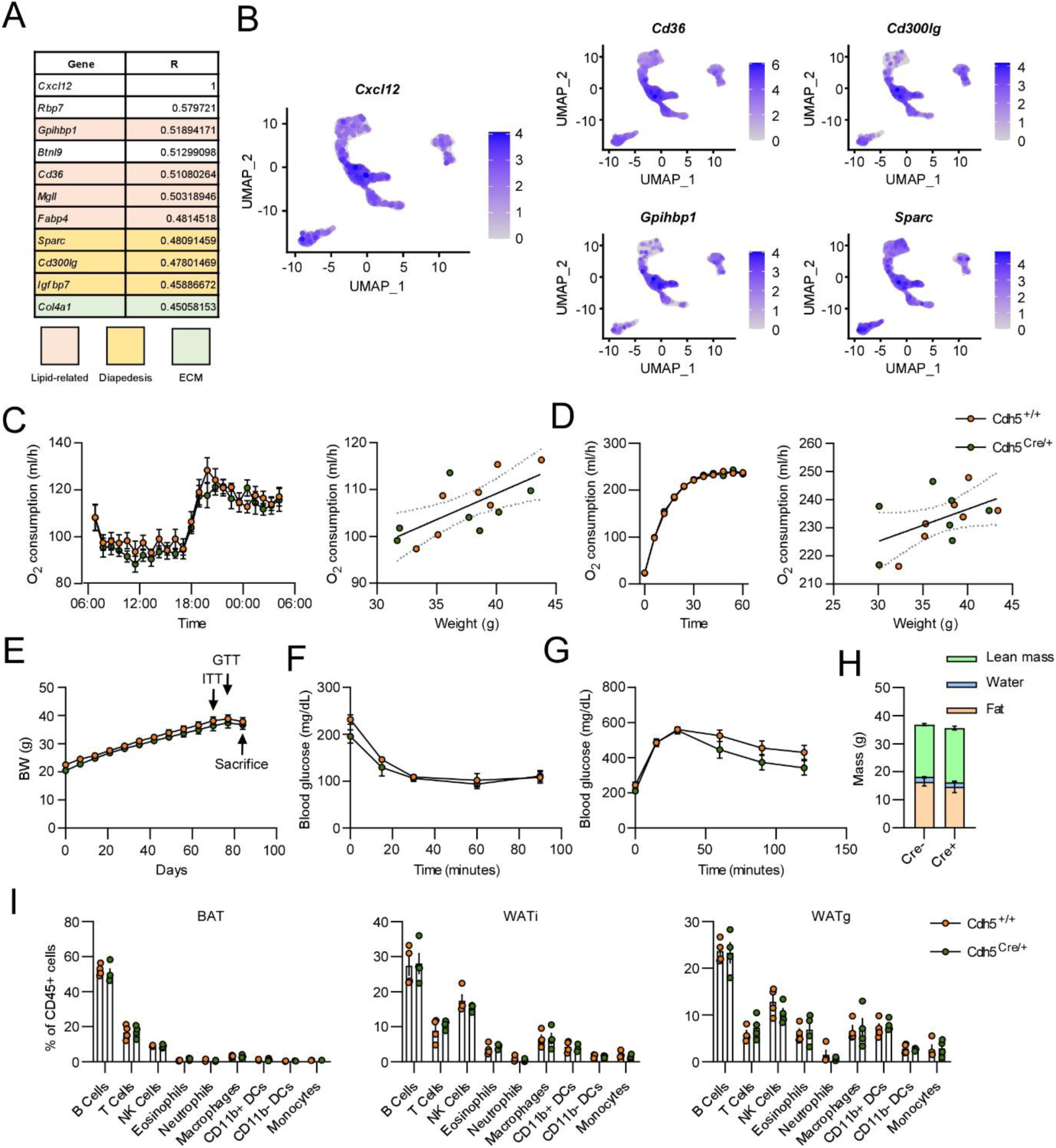
Endothelial CXCL12 is dispensable for adipose tissue function under obesogenic conditions. (**A**) Single-cell gene expression correlation of *Cxcl12* and lipid- and immune cell diapedesis-related genes in WATi-ECs. (**B**) UMAP plots showing similar gene expression patterns of *Cxcl12*, *Cd36*, *Gpihbp1*, *Cd300lg*, and *Sparc* in WATi-ECs. (**C**) O_2_ consumption of CXCL12-ECKO mice and WT littermates after 12 weeks of HFD feeding at RT (ANCOVA). (**D**) O_2_ consumption of HFD-fed CXCL12 mice and WT littermates during a 1-hour cold (4° C) challenge (ANCOVA). (**E**) Body weight (BW) of HFD-fed CXCL12-ECKO mice and WT littermates. (**F**) Insulin tolerance test in HFD-fed CXCL12-ECKO mice and WT littermates. (**G**) Glucose tolerance test in HFD-fed CXCL12-ECKO mice and WT littermates. (**H**) Body composition measured by NMR of HFD-fed CXCL12-ECKO mice and WT littermates. (**I**) FACS analysis of immune cell contents in BAT, WATi, and WATg of HFD-fed CXCL12-ECKO mice and WT littermates.

To investigate a potential role for EC-derived CXCL12 in adipose tissue inflammation, we performed a comprehensive FACS-based analysis of immune cell contents of all three major adipose tissues (BAT, WATi, and WATg) in HFD-fed CXCL12-ECKO mice and WT littermates. Surprisingly, we found no differences in B-cells, T-cells, natural killer (NK) cells, eosinophils, neutrophils, macrophages, dendritic cells, or monocytes (Figure 6I). Additionally, analysis of macrophage subsets revealed an unchanged proportion of the different ATM populations in mice with endothelial-specific CXCL12-KO compared to littermate controls. (Supplemental Figure 3F). Histological analysis of the different adipose tissue depots did not reveal any changes in morphology (Supplemental Figure 3G).

Taken together, these data show that EC-derived CXCL12 is dispensable for lipid storage and adipose tissue inflammation in obesogenic conditions.

## 4. Discussion

CXCL12 is thought to be one of the first homeostatic chemokines to evolve in vertebrates [36], has an essential role in embryonic development, and is expressed constitutively in many organs in adulthood. In its role as a homeostatic chemokine, CXCL12 regulates several key cellular processes, such as immune cell trafficking, angiogenesis, neurogenesis, and stemness [10–12,37]. These processes are of critical importance in adipose tissue remodeling under various conditions, e.g. cold adaptation and obesity.

In our study, we found that ECs are the dominant source of CXCL12 in BAT, and that the expression of *Cxcl12* in BAT ECs is highly upregulated during cold acclimation. Therefore, we hypothesized that CXCL12 generated by ECs, the cell type with the highest *Cxcl12* expression in adipose tissue, could be an important player in adipose tissue biology. Previous studies have shown that smooth muscle-derived CXCL12 influences adipose tissue function by regulating influx of CD301^+^ macrophages that in turn support neuronal maintenance [23]. Other studies have suggested that mature brown adipocytes are a source CXCL12 that might act either as an immune cell chemoattractant, a direct activator of neurogenesis, or an autocrine activator of the brown adipocytes themselves [24]. Interestingly, our detailed and stringent FACS-based analysis of the AT-EC transcriptome, as well as re-analysis of single-cell sequencing data, showed that ECs are the major source of CXCL12 in BAT. We could also show that the “floating fraction” of collagenase-digested BAT, which is often assumed to only contain mature, lipid-laden cells, appears to contain a significant amount of ECs, possibly owing to high cell-cell adhesion between ECs and brown adipocytes, which has also been suggested by previous studies [31]. Importantly, whole BAT tissue extracts from CXCL12-ECKO mice showed extremely low levels of *Cxcl12* expression, further indicating that ECs are the major source of CXCL12 in BAT, especially during cold exposure.

Our *in vitro* experiments performed on muMEC microvascular ECs as well as primary BAT SVF spheroids showed that CXCL12 significantly enhances microvascular angiogenesis and vascularization. Additionally, treatment of primary sympathetic neurons with CXCL12 increased neurite outgrowth and branching. Given these results, we set out to identify the functional role of EC-derived CXCL12 in adipose tissues, with a special focus on BAT. To our surprise, we found that ablation of CXCL12 in ECs had no significant impact on any of the major functions of adipose tissues. In mice housed at room temperature, knock-down of Cxcl12 did not affect either energy expenditure, body- and AT weight, food intake, or motility. Additionally, we found no evidence of altered vascularization, expression of thermogenic markers, or tissue morphology. Mice subjected to a one-week cold challenge also did not display any metabolic phenotype, with normal energy expenditure, body weight, food intake, motility, BAT innervation, and thermogenic marker expression. Under obesogenic conditions (DIO), we found that CXCL12-ECKO mice had normal lipid storage capacity, cold tolerance, glucose handling, and inflammatory response.

A limitation of this study is the use of a non-inducible Cre to knock out *Cxcl12* in ECs, which did not allow for temporal control over the *Cxcl12* deletion. This could potentially lead to compensation for the lack of CXCL12 on either vascularization or neurogenesis by other mechanisms and pathways. However, the lack of phenotype observed in the CXCL12-ECKO mice more likely suggests that endothelial *Cxcl12* expression is dispensable for both adipose tissue development as well as function in adult mice. Additionally, it should be noted that, although *Cxcl12* is more highly expressed in ECs, single cell sequencing data also show expression of *Cxcl12* in preadipocytes, macrophages, and to some extent smooth muscle cells. Given the close proximity of both preadipocytes and smooth muscle cells to the endothelium, it is conceivable that the lack of *Cxcl12* expression in ECs could be compensated by CXCL12 derived from these other cell types in adipose tissues.

CXCL12 has a plethora of effects on various cell types under different stimuli. Our *in vitro* studies showed a clear effect of CXCL12 on angiogenesis and neurite outgrowth in native BAT ECs and primary sympathetic neurons. In addition to this, CXCL12 could potentially also regulate other processes that are relevant for adipose tissue function, e.g. preadipocyte stemness and vascular permeability. Although we did not directly investigate these parameters, it seems clear from our data that any effect of EC-derived CXCL12 on e.g. preadipocytes or the ECs themselves are not essential for adipose tissue function either during cold stress or under obesogenic conditions. Other paracrine functions of EC-derived CXCL12 could potentially be the regulation of smaller subsets of immune cells that were not distinguishable *via* our FACS panel. Sensory nerves may also express *Ackr3* or *Cxcr4* and thereby be regulated by EC-derived CXCL12. Overall, our *in vivo* studies clearly demonstrate that EC-derived CXCL12 does not influence these processes in a functionally relevant manner during either cold stress or DIO.

Endothelial CXCL12 has previously been shown to play important roles in various physiological and adaptive processes in different organs. In the lung, EC-derived CXCL12 defines endothelial niches and promotes vascularization during development [38]. In the brain, endothelial CXCL12 is responsible for attracting NK cells during ischemic stroke [39]. Additionally, in addition to the previously mentioned effect on *in vitro* HUVEC angiogenesis [28], CXCL12-CXCR4 signaling has been shown to regulate tumor angiogenesis *in vivo* [40]. It is puzzling that, in AT, EC-derived CXCL12 is dispensable for tissue function, given the high expression of *Cxcl12* in these cells in AT. However, it is worth noting that chemokine signaling often displays significant redundancy, and it is therefore possible that other chemokines, e.g. CXCL11, can sufficiently compensate for the lack of CXCL12 in our mouse model. Alternatively, CXCL12 derived from other perivascular cell types, e.g. mesenchymal stem cells, smooth muscle cells, or pericytes, might be able to support proper AT function even in the absence of endothelial CXCL12.

## 5. Conclusions

In conclusion, our study shows that endothelial CXCL12 is dispensable for normal adipose tissue function. Given the previously described roles of CXCL12 in adipose tissues, together with the high expression of *Cxcl12* in adipose tissue ECs, this finding is surprising. Although we subjected CXCL12-ECKO mice to both metabolic and thermogenic challenges, we could not detect a functional role for endothelium-derived CXCL12 under these conditions.

## Supplementary Materials

Supplemental Figure 1: Gating strategy for FACS-isolation of adipose tissue ECs; Supplemental Figure 2: Characterization of CXCL12-ECKO (Cdh5^Cre/+^) mice under normal housing conditions; Supplemental Figure 3: Endothelial CXCL12 is dispensable for adipose tissue function under obesogenic conditions; Full uncropped Western blots for Figure 4F.

## Author Contributions

S.H.: Conceptualization, Data Curation, Formal Analysis, Investigation, Methodology, Project Administration, Supervision, Visualization, Writing – Original Draft Preparation, Writing – Review & Editing. M.R.: Investigation, Formal Analysis. B.E.F.P.: Investigation, Formal Analysis. F.J.G.: Investigation, Formal Analysis. S.L.: Investigation, Formal Analysis. I.F.: Supervision, Funding Acquisition, Writing – Review & Editing. A.P.: Conceptualization, Funding Acquisition, Resources, Project Administration, Supervision, Writing – Review & Editing. All authors contributed to the final manuscript.

## Institutional Review Board Statement

The animal study protocol was approved by the Landesamt für Verbraucherschutz und Ernährung, NRW, Germany, with the approval numbers 81-02.04.2023.A113 and 81-02.04.2015.A202.

## Data Availability Statement

All data will be made available upon reasonable request.

## Acknowledgments

The authors would like to thank Hannah Lamby and Markus Raspe for technical assistance. We would also like to thank the Bioinformatics and Next Generation Sequencing Core Facilities of the Medical Faculty at the University of Bonn. Parts of some figures were made with BioRender (https://www.biorender.com/).

## Abbreviations

The following abbreviations are used in this manuscript:

AT: Adipose tissue
BAT: Brown adipose tissue
EC: Endothelial cell
WAT: White adipose tissue
CXCL12 C-X-C: motif chemokine 12
ATEC: Adipose tissue endothelial cell
WT: Wild-type
HFD: High fat diet
BABB: Benzyl alcohol and benzyl benzoate
WATi: Inguinal white adipose tissue
WATg: Gonadal white adipose tissue
FACS: Fluorescence-Activated Cell Sorting
MACS: Magnetic-activated cell sorting
HUVEC: Human umbilical cord vein endothelial cells
ECKO: Endothelial cell knock-out
ECM: Extracellular matrix
TH: Tyrosine hydroxylase
RT: Room temperature
DIO: Diet-induced obesity
NK: Natural Killer

## References

1. Hildebrand, S.; Pfeifer, A. The obesity pandemic and its impact on non-communicable disease burden. Pflugers Arch 2025, 477, 657–668, doi:10.1007/s00424-025-03066-8.

2. Pfeifer, A.; Hoffmann, L.S. Brown, beige, and white: the new color code of fat and its pharmacological implications. Annu Rev Pharmacol Toxicol 2015, 55, 207–227, doi:10.1146/annurev-pharmtox-010814-124346.

3. van Marken Lichtenbelt, W.D.; Vanhommerig, J.W.; Smulders, N.M.; Drossaerts, J.M.; Kemerink, G.J.; Bouvy, N.D.; Schrauwen, P.; Teule, G.J. Cold-activated brown adipose tissue in healthy men. N Engl J Med 2009, 360, 1500–1508, doi:10.1056/NEJMoa0808718.

4. Saito, M.; Okamatsu-Ogura, Y.; Matsushita, M.; Watanabe, K.; Yoneshiro, T.; Nio-Kobayashi, J.; Iwanaga, T.; Miyagawa, M.; Kameya, T.; Nakada, K.; et al. High incidence of metabolically active brown adipose tissue in healthy adult humans: effects of cold exposure and adiposity. Diabetes 2009, 58, 1526–1531, doi:10.2337/db09-0530.

5. Cypess, A.M.; Lehman, S.; Williams, G.; Tal, I.; Rodman, D.; Goldfine, A.B.; Kuo, F.C.; Palmer, E.L.; Tseng, Y.H.; Doria, A.; et al. Identification and importance of brown adipose tissue in adult humans. N Engl J Med 2009, 360, 1509–1517, doi:10.1056/NEJMoa0810780.

6. Becher, T.; Palanisamy, S.; Kramer, D.J.; Eljalby, M.; Marx, S.J.; Wibmer, A.G.; Butler, S.D.; Jiang, C.S.; Vaughan, R.; Schoder, H.; et al. Brown adipose tissue is associated with cardiometabolic health. Nat Med 2021, 27, 58–65, doi:10.1038/s41591-020-1126-7.

7. Jo, J.; Gavrilova, O.; Pack, S.; Jou, W.; Mullen, S.; Sumner, A.E.; Cushman, S.W.; Periwal, V. Hypertrophy and/or Hyperplasia: Dynamics of Adipose Tissue Growth. PLoS Comput Biol 2009, 5, e1000324, doi:10.1371/journal.pcbi.1000324.

8. Alkhouri, N.; Gornicka, A.; Berk, M.P.; Thapaliya, S.; Dixon, L.J.; Kashyap, S.; Schauer, P.R.; Feldstein, A.E. Adipocyte apoptosis, a link between obesity, insulin resistance, and hepatic steatosis. J Biol Chem 2010, 285, 3428–3438, doi:10.1074/jbc.M109.074252.

9. Bartelt, A.; Heeren, J. Adipose tissue browning and metabolic health. Nat Rev Endocrinol 2014, 10, 24–36, doi:10.1038/nrendo.2013.204.

10. Surmi, B.K.; Hasty, A.H. Macrophage infiltration into adipose tissue: initiation, propagation and remodeling. Future Lipidol 2008, 3, 545–556, doi:10.2217/17460875.3.5.545.

11. Herold, J.; Kalucka, J. Angiogenesis in Adipose Tissue: The Interplay Between Adipose and Endothelial Cells. Front Physiol 2020, 11, 624903, doi:10.3389/fphys.2020.624903.

12. Willows, J.W.; Blaszkiewicz, M.; Townsend, K.L. The Sympathetic Innervation of Adipose Tissues: Regulation, Functions, and Plasticity. Compr Physiol 2023, 13, 4985–5021, doi:10.1002/cphy.c220030.

13. Fischer, A.W.; Jaeckstein, M.Y.; Gottschling, K.; Heine, M.; Sass, F.; Mangels, N.; Schlein, C.; Worthmann, A.; Bruns, O.T.; Yuan, Y.; et al. Lysosomal lipoprotein processing in endothelial cells stimulates adipose tissue thermogenic adaptation. Cell Metab 2021, 33, 547–564 e547, doi:10.1016/j.cmet.2020.12.001.

14. Sengenes, C.; Miranville, A.; Lolmede, K.; Curat, C.A.; Bouloumie, A. The role of endothelial cells in inflamed adipose tissue. J Intern Med 2007, 262, 415–421, doi:10.1111/j.1365-2796.2007.01853.x.

15. Daquinag, A.C.; Gao, Z.; Yu, Y.; Kolonin, M.G. Endothelial TrkA coordinates vascularization and innervation in thermogenic adipose tissue and can be targeted to control metabolism. Mol Metab 2022, 63, 101544, doi:10.1016/j.molmet.2022.101544.

16. Shi, Y.; Huang, X.; Zeng, Y.; Zhai, M.; Yao, H.; Liu, C.; Li, B.; Gong, S.; Yu, Q.; Zhuang, J.; et al. Endothelial TET2 regulates the white adipose browning and metabolism via fatty acid oxidation in obesity. Redox Biol 2024, 69, 103013, doi:10.1016/j.redox.2023.103013.

17. Speyer, C.L.; Ward, P.A. Role of endothelial chemokines and their receptors during inflammation. J Invest Surg 2011, 24, 18–27, doi:10.3109/08941939.2010.521232.

18. van Gemst, J.J.; Kouwenberg, M.; Rops, A.; van Kuppevelt, T.H.; Berden, J.H.; Rabelink, T.J.; Loeven, M.A.; van der Vlag, J. Differential binding of chemokines CXCL1, CXCL2 and CCL2 to mouse glomerular endothelial cells reveals specificity for distinct heparan sulfate domains. PLoS One 2018, *13*, e0201560, doi:10.1371/journal.pone.0201560.

19. Xue, Y.; Petrovic, N.; Cao, R.; Larsson, O.; Lim, S.; Chen, S.; Feldmann, H.M.; Liang, Z.; Zhu, Z.; Nedergaard, J.; et al. Hypoxia-independent angiogenesis in adipose tissues during cold acclimation. Cell Metab 2009, 9, 99–109, doi:10.1016/j.cmet.2008.11.009.

20. Wang, Y.; Leung, V.H.; Zhang, Y.; Nudell, V.S.; Loud, M.; Servin-Vences, M.R.; Yang, D.; Wang, K.; Moya-Garzon, M.D.; Li, V.L.; et al. The role of somatosensory innervation of adipose tissues. Nature 2022, 609, 569–574, doi:10.1038/s41586-022-05137-7.

21. Grasman, J.M.; Kaplan, D.L. Human endothelial cells secrete neurotropic factors to direct axonal growth of peripheral nerves. Sci Rep 2017, 7, 4092, doi:10.1038/s41598-017-04460-8.

22. Bajoghli, B. Evolution and function of chemokine receptors in the immune system of lower vertebrates. Eur J Immunol 2013, 43, 1686–1692, doi:10.1002/eji.201343557.

23. Lee, D.; Benvie, A.M.; Steiner, B.M.; Kolba, N.J.; Ford, J.G.; McCabe, S.M.; Jiang, Y.; Berry, D.C. Smooth muscle cell-derived Cxcl12 directs macrophage accrual and sympathetic innervation to control thermogenic adipose tissue. Cell Rep 2024, 43, 114169, doi:10.1016/j.celrep.2024.114169.

24. Agueda-Oyarzabal, M.; Isidor, M.S.; Plucinska, K.; Ingerslev, L.R.; Dmytriyeva, O.; Petersen, P.S.S.; Laftih, S.; Pontoppidan, A.B.; Henningsen, J.B.; Rupar, K.; et al. Transcriptomic signatures of cold acclimated adipocytes reveal CXCL12 as a Brown autocrine and paracrine chemokine. Mol Metab 2025, 93, 102102, doi:10.1016/j.molmet.2025.102102.

25. Kim, D.; Kim, J.; Yoon, J.H.; Ghim, J.; Yea, K.; Song, P.; Park, S.; Lee, A.; Hong, C.P.; Jang, M.S.; et al. CXCL12 secreted from adipose tissue recruits macrophages and induces insulin resistance in mice. Diabetologia 2014, 57, 1456–1465, doi:10.1007/s00125-014-3237-5.

26. Kurita, K.; Ishikawa, K.; Takeda, K.; Fujimoto, M.; Ono, H.; Kumagai, J.; Inoue, H.; Yokoh, H.; Yokote, K. CXCL12-CXCR4 pathway activates brown adipocytes and induces insulin resistance in CXCR4-deficient mice under high-fat diet. Sci Rep 2019, 9, 6165, doi:10.1038/s41598-019-42127-8.

27. Shamsi, F.; Piper, M.; Ho, L.L.; Huang, T.L.; Gupta, A.; Streets, A.; Lynes, M.D.; Tseng, Y.H. Vascular smooth muscle-derived Trpv1(+) progenitors are a source of cold-induced thermogenic adipocytes. Nat Metab 2021, 3, 485–495, doi:10.1038/s42255-021-00373-z.

28. Zhang, M.; Qiu, L.; Zhang, Y.; Xu, D.; Zheng, J.C.; Jiang, L. CXCL12 enhances angiogenesis through CXCR7 activation in human umbilical vein endothelial cells. Sci Rep 2017, 7, 8289, doi:10.1038/s41598-017-08840-y.

29. Stumm, R.K.; Zhou, C.; Ara, T.; Lazarini, F.; Dubois-Dalcq, M.; Nagasawa, T.; Hollt, V.; Schulz, S. CXCR4 regulates interneuron migration in the developing neocortex. J Neurosci 2003, 23, 5123–5130, doi:10.1523/JNEUROSCI.23-12-05123.2003.

30. Lopez-Bendito, G.; Sanchez-Alcaniz, J.A.; Pla, R.; Borrell, V.; Pico, E.; Valdeolmillos, M.; Marin, O. Chemokine signaling controls intracortical migration and final distribution of GABAergic interneurons. J Neurosci 2008, 28, 1613–1624, doi:10.1523/JNEUROSCI.4651-07.2008.

31. Wang, T.; Sharma, A.K.; Wolfrum, C. Novel insights into adipose tissue heterogeneity. Rev Endocr Metab Disord 2022, 23, 5–12, doi:10.1007/s11154-021-09703-8.

32. Kalucka, J.; de Rooij, L.; Goveia, J.; Rohlenova, K.; Dumas, S.J.; Meta, E.; Conchinha, N.V.; Taverna, F.; Teuwen, L.A.; Veys, K.; et al. Single-Cell Transcriptome Atlas of Murine Endothelial Cells. Cell 2020, 180, 764–779 e720, doi:10.1016/j.cell.2020.01.015.

33. Dotan, I.; Werner, L.; Vigodman, S.; Weiss, S.; Brazowski, E.; Maharshak, N.; Chen, O.; Tulchinsky, H.; Halpern, Z.; Guzner-Gur, H. CXCL12 is a constitutive and inflammatory chemokine in the intestinal immune system. Inflamm Bowel Dis 2010, 16, 583–592, doi:10.1002/ibd.21106.

34. Panitz, N.; Theisgen, S.; Samsonov, S.A.; Gehrcke, J.P.; Baumann, L.; Bellmann-Sickert, K.; Kohling, S.; Pisabarro, M.T.; Rademann, J.; Huster, D.; et al. The structural investigation of glycosaminoglycan binding to CXCL12 displays distinct interaction sites. Glycobiology 2016, 26, 1209–1221, doi:10.1093/glycob/cww059.

35. Lantermans, H.C.; Ma, F.; Kuil, A.; de Rooij, M.F.M.; Bergkamp, N.D.; van der Meer, W.; van Buul, J.D.; Smit, M.J.; Kersten, M.J.; Spaargaren, M.; et al. Presentation of CXCL12gamma by heparan sulfate proteoglycans activates CXCR4 without desensitization in normal and malignant B cells. Blood Adv 2025, 9, 2307–2320, doi:10.1182/bloodadvances.2024014396.

36. Nomiyama, H.; Osada, N.; Yoshie, O. Systematic classification of vertebrate chemokines based on conserved synteny and evolutionary history. Genes Cells 2013, 18, 1–16, doi:10.1111/gtc.12013.

37. Singh, D.D.; Yadav, D.K.; Shin, D. Targeting the CXCR4/CXCL12 Axis to Overcome Drug Resistance in Triple-Negative Breast Cancer. Cells 2025, 14, doi:10.3390/cells14181482.

38. Chandrasekaran, P.; Negretti, N.M.; Sivakumar, A.; Liberti, D.C.; Wen, H.; Peers de Nieuwburgh, M.; Wang, J.Y.; Michki, S.N.; Chaudhry, F.N.; Kaur, S.;, et al. CXCL12 defines lung endothelial heterogeneity and promotes distal vascular growth. Development 2022, 149, doi:10.1242/dev.200909.

39. Huang, C.L.; Huang, Y.N.; Yao, L.; Li, J.P.; Zhang, Z.H.; Huang, Z.Q.; Chen, S.X.; Zhang, Y.L.; Wang, J.F.; Chen, Y.X.; et al. Thoracic perivascular adipose tissue inhibits VSMC apoptosis and aortic aneurysm formation in mice via the secretome of browning adipocytes. Acta Pharmacol Sin 2023, 44, 345–355, doi:10.1038/s41401-022-00959-7.

40. Liang, Z.; Brooks, J.; Willard, M.; Liang, K.; Yoon, Y.; Kang, S.; Shim, H. CXCR4/CXCL12 axis promotes VEGF-mediated tumor angiogenesis through Akt signaling pathway. Biochem Biophys Res Commun 2007, 359, 716–722, doi:10.1016/j.bbrc.2007.05.182.

