## Supplemental Data for "Endothelial CXCL12 is dispensable for adipose tissue function"

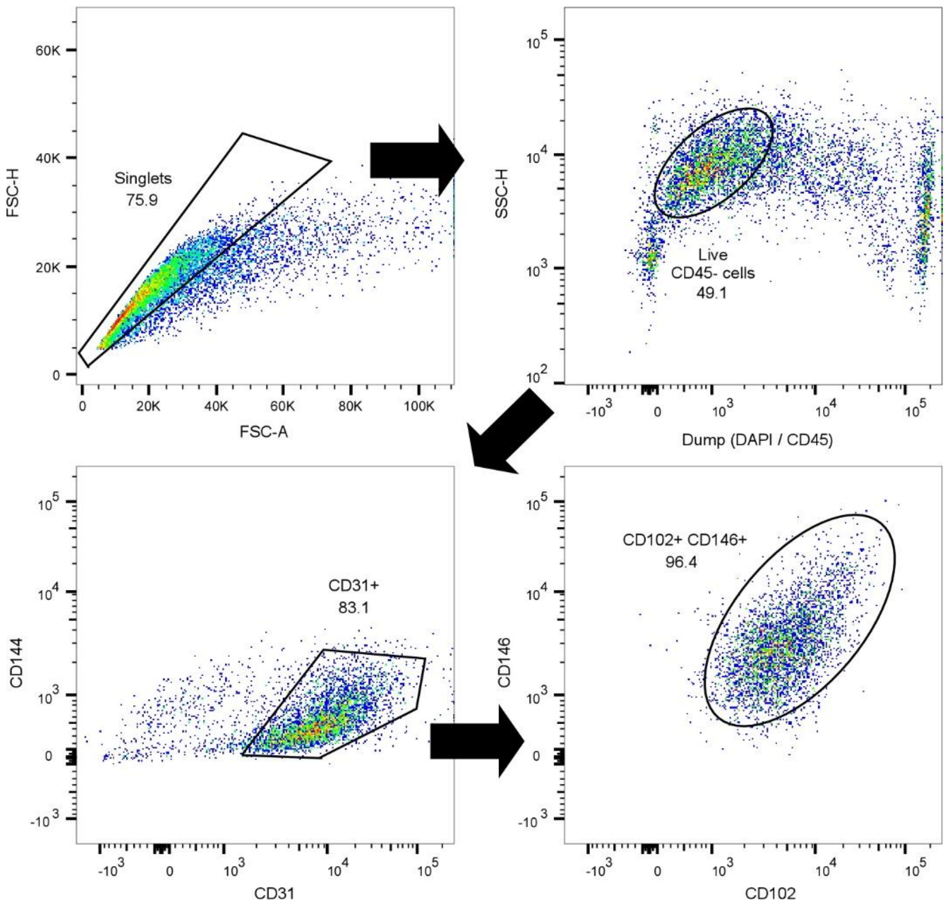


**Supplemental Figure 1.** Gating strategy for FACS-isolation of adipose tissue ECs.

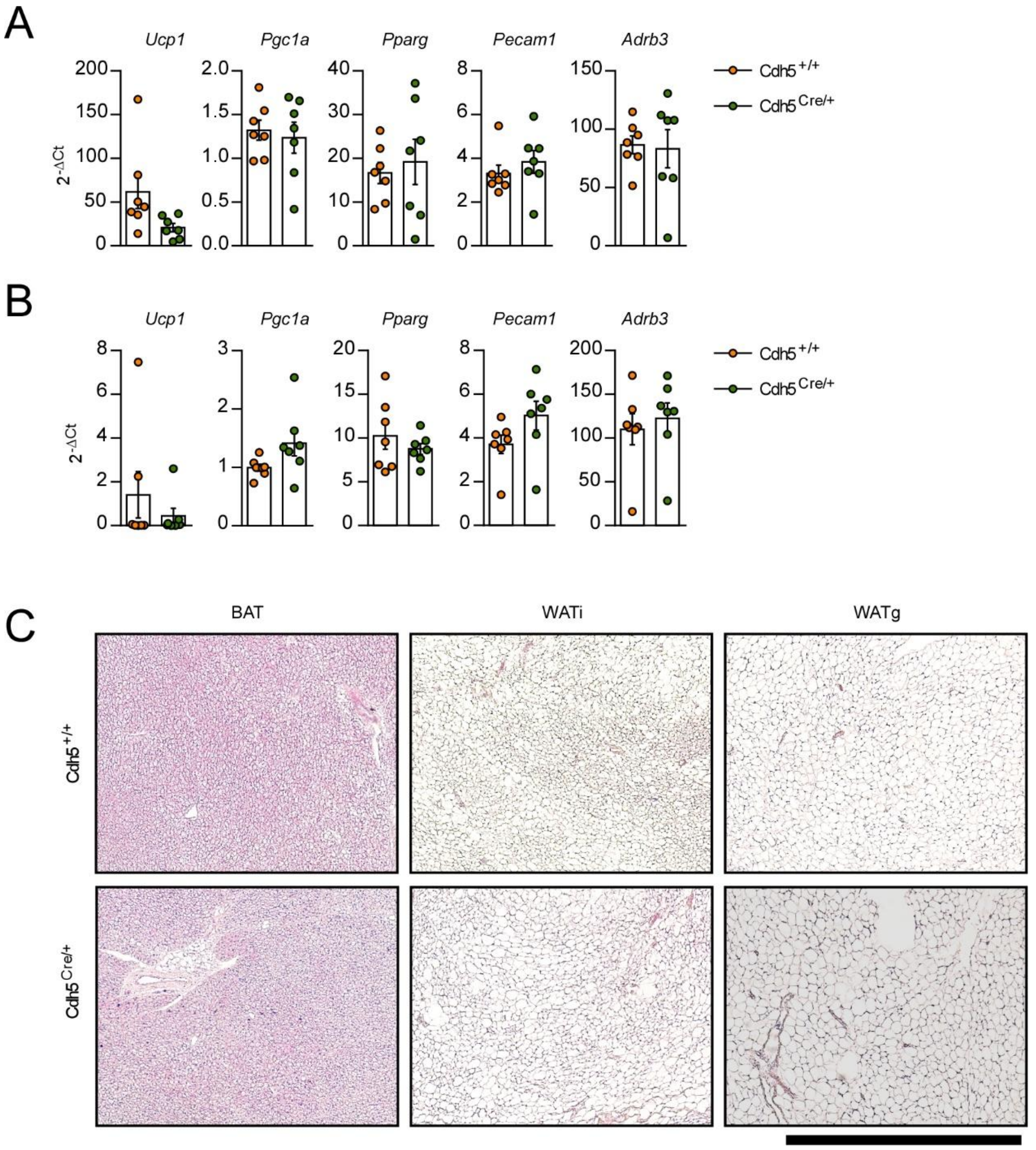


**Supplemental Figure 2.** Characterization of CXCL12-ECKO (Cdh5$^{Cre/+}$) mice under normal housing conditions. (**A**) mRNA expression of *Ucp1*, *Pgc1a*, *Pparg*, *Pecam1,* and *Adrb3* in the WATi of CXCL12-ECKO mice and WT littermates. (**B**) mRNA expression of *Ucp1*, *Pgc1a*, *Pparg*, *Pecam1,* and *Adrb3* in the WATg of CXCL12-ECKO mice and WT littermates. (**C**) H&E staining of histological sections from BAT, WATi, and WATg of CXCL12-ECKO mice and WT littermates. Scale bar = 1 mm.

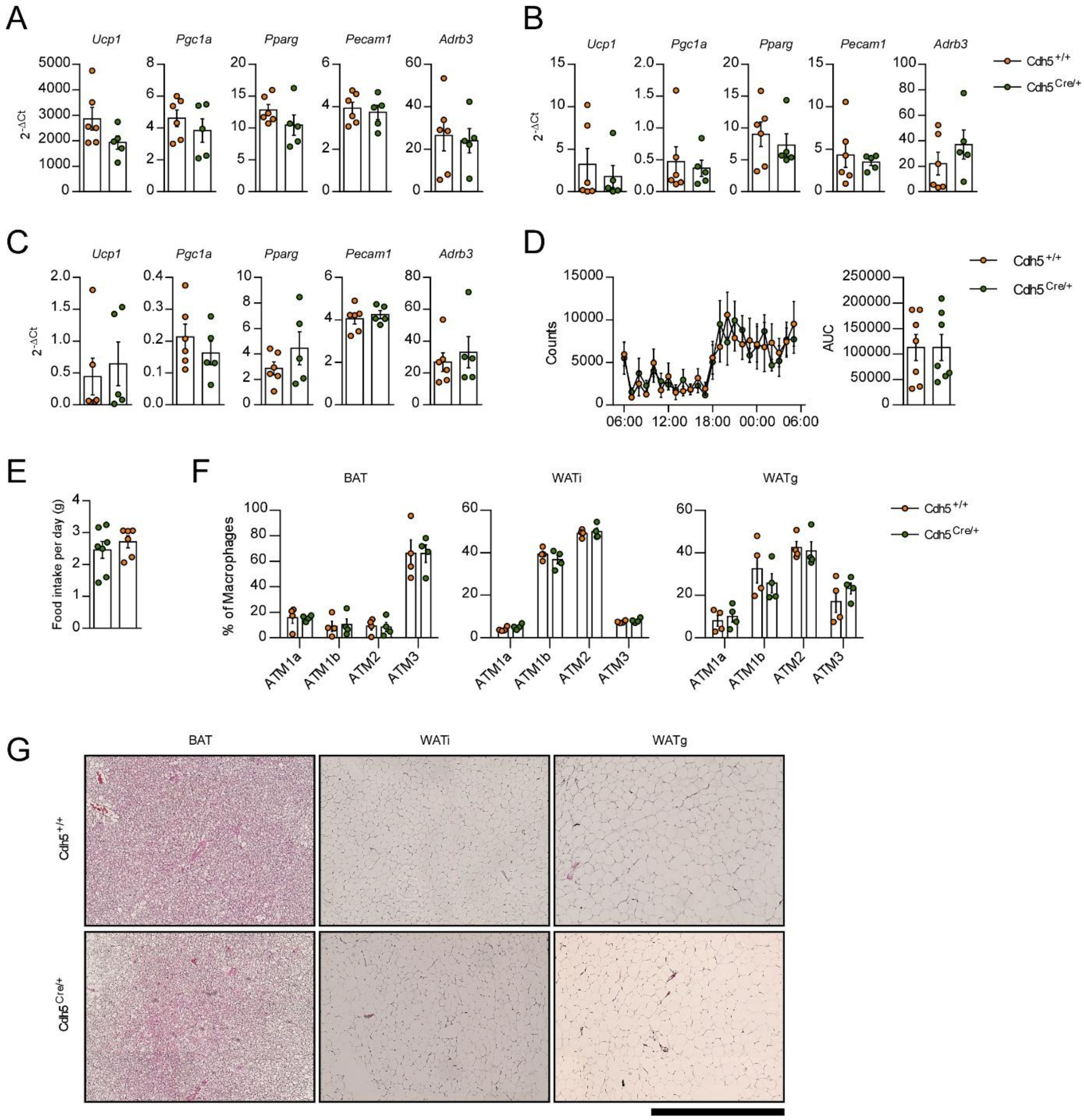


**Supplemental Figure 3:** Endothelial CXCL12 is dispensable for adipose tissue function under obesogenic conditions. (**A**) mRNA expression of *Ucp1*, *Pgc1a*, *Pparg*, *Pecam1*, and *Adrb3* in BAT of HFD-fed CXCL12-ECKO mice and WT littermates. (**B**) mRNA expression of *Ucp1*, *Pgc1a*, *Pparg*, *Pecam1*, and *Adrb3* in WATi of HFD-fed CXCL12-ECKO mice and WT littermates. (**C**) mRNA expression of *Ucp1*, *Pgc1a*, *Pparg*, *Pecam1*, and *Adrb3* in WATg of HFD-fed CXCL12-ECKO mice and WT littermates. (**D**) 24-hour motility of HFD-fed CXCL12-ECKO mice and WT littermates housed at 23° C. (**E**) Daily food consumption of HFD-fed CXCL12-ECKO mice and

WT littermates. (**F**) Relative abundance of macrophage subsets in adipose tissues of HFD-fed CXCL12-ECKO mice and WT littermates. (**G**) H&E staining of histological sections from BAT, WATi, and WATg of HFD-fed CXCL12-ECKO mice and WT littermates. Scale bar = 1 mm.

Full uncropped blots for Figure 4F:

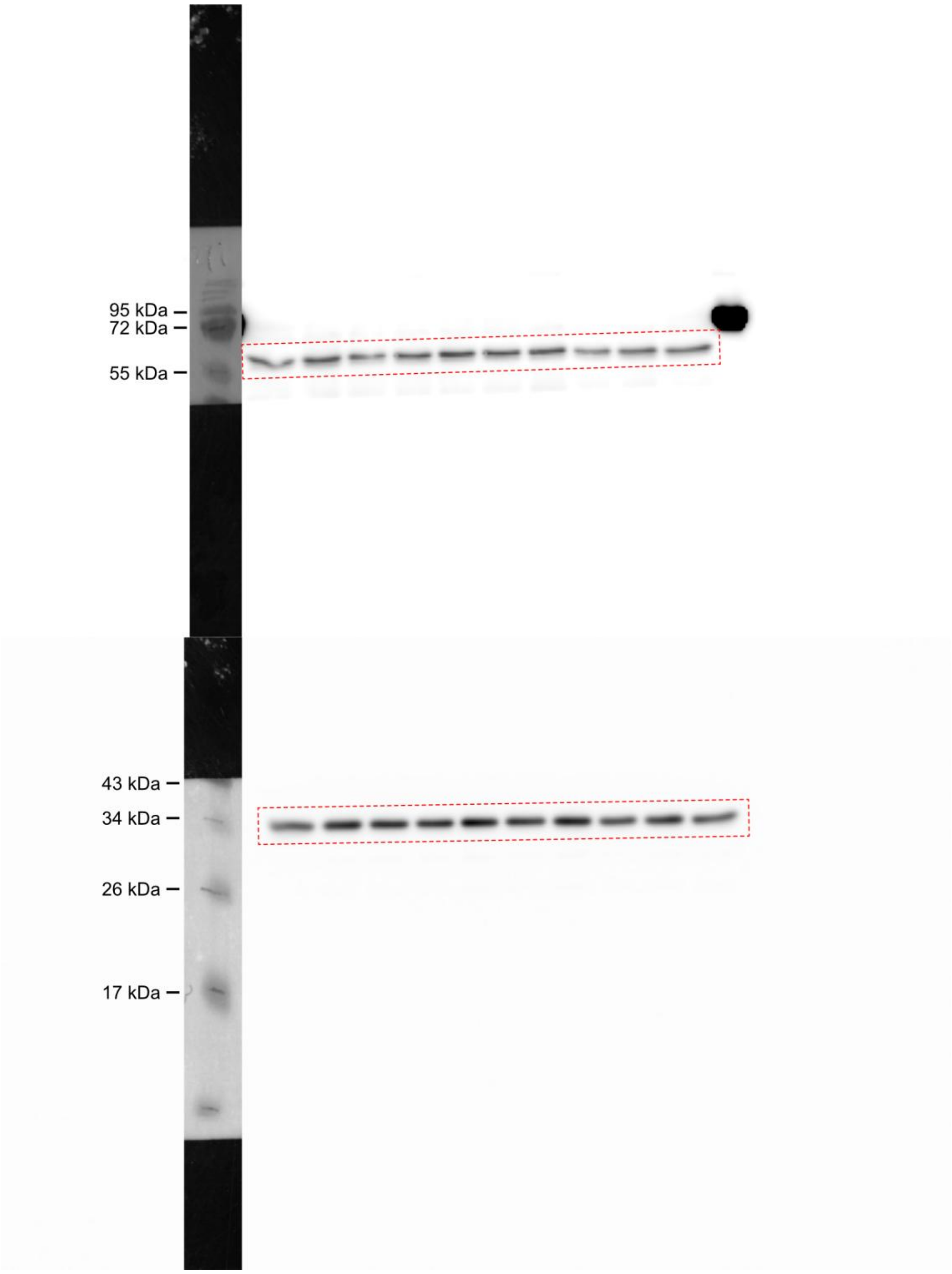
